# Human neutrophils communicate remotely via glutamate-induced glutamate release

**DOI:** 10.1101/2022.02.25.482046

**Authors:** Olga Kopach, Sergyi Sylantyev, Lucie Bard, Piotr Michaluk, Ana Gutierrez del Arroyo, Gareth L. Ackland, Alexander V. Gourine, Dmitri A. Rusakov

## Abstract

Neutrophils are white blood cells that are critical to the acute inflammatory and adaptive immune responses. Their swarming-pattern behaviour is controlled by multiple cellular cascades involving calcium-dependent release of various signalling molecules. Previous studies have reported that neutrophils express glutamate receptors and can release glutamate but evidence of direct neutrophil-neutrophil communication has been elusive. Here, we hold semi-suspended cultured human neutrophils in patch-clamp whole-cell mode to find that calcium mobilisation induced by stimulating one neutrophil can trigger an NMDA receptor-driven membrane current and calcium signal in neighbouring neutrophils. We employ an enzymatic-based imaging assay to image, in real time, glutamate release from neutrophils induced by glutamate released from their neighbours. These observations provide direct evidence for a positive-feedback inter-neutrophil communication that could contribute to mechanisms regulating communal neutrophil behaviour.

## Introduction

Neutrophils provide the first line of defence against pathogens during acute inflammation ^1–3^. In humans, a significant decrease in neutrophil numbers could lead to severe immunodeficiency or death. These polymorphonuclear leukocytes mobilise to the site of inflammation and engage several cell-killing mechanisms to clear the infection ^4,5^. Cellular mechanisms underpinning communication of neutrophils with other actors of a pathological immune response, such as platelets, T-cells, or infectious agents, have been intensely studied ^6–8^.

Nonetheless important is the rapidly emerging knowledge of intercellular communication among neutrophils themselves. Cooperative, swarm-like migration patterns of neutrophils have been considered an essential process in their tissue response ^9,10^. The underpinning molecular mechanisms involve the lipid LTB4 and integrins, the release of signalling molecules such as ATP ^11^, and the action of connexins accompanied by cooperative calcium alarm signals ^12^. Calcium-dependent release of chemo-attractants enables self-sustained paracrine signalling, thus providing positive-feedback amplification which drives self-organised neutrophil ensemble behaviour ^13–15^. Generated in a small group of clustering neutrophils, the molecular signal thus triggers what appears to be a chain-reaction mechanism communicating with more distant cells attracting them for further swarm growth ^15^. However, direct evidence for positive-feedback neutrophil-neutrophil communication at the cellular level has been scarce.

Ca^2+^ signalling has long been considered key to the physiological functions of neutrophils ^16^, which are equipped with a variety of cell-surface receptors, including GPCRs, FcRs, and integrins, capable of mediating Ca^2+^ messages ^17–19^. Previous work has shown that neutrophils can secrete glutamate and D-serine ^20,21^ and express NMDA receptors (NMDARs) ^21^. Because NMDAR activation by glutamate and the co-agonist D-serine can generate major Ca^2+^ influx and thus engage release machinery in the host cell, we thought it was important to understand whether glutamate-induced glutamate release could contribute to the positive-feedback signal amplification among neutrophil ensembles. To explore this, we engaged up-to-date neuroscience techniques that we previously established while exploring among central neurons and astrocytes ^22–25^.

## STUDY DESIGN

A detailed description of the Methods is supplied in the Supplementary Data (available on the Blood website). In brief, neutrophils were isolated from blood samples obtained by venepuncture from healthy volunteers according to the protocols approved by the UCL Research Ethics Committee (UCL Queen Square Institute of Neurology, London, UK), with the informed consent of participants. We used a method of dextran sedimentation and differential centrifugation through a Ficoll-Hypaque density gradient ^26^. Residual erythrocytes were removed by hypotonic lysis; neutrophils were plated on coverslips coated with poly-DL-Lysine and laminine and were maintained at 37°C (95% O_2_ / 5% CO_2_) until used. The experiment presented here were carried out in freshly isolated neutrophils (Supplemental Methods); healthy cells were identified by their normal morphology, microscopic motility (such as ’attacking’ the patch pipette tip with pseudopodia), normal membrane potential in whole-cell, and reversible Ca^2+^ signalling.

Visualised patch-clamp recordings from the neutrophils were performed as described earlier for small cerebellar neurons ^27^, and detailed in Supplemental Methods. Neutrophils were held whole-cell in either current-clamp or voltage-clamp mode, as indicated. The previously established piezo-driven theta-glass fast-application system ^23^ was employed to probe NMDA receptor sensitivity of individual neutrophils.

For Ca^2+^ imaging, neutrophils were loaded with the Ca^2+^-indicator Fluo-4/AM (5 μM). Two-photon excitation imaging was carried out using an Olympus FV-1000MPE or Femtonics Femto-2D system, as detailed previously ^28,29^ and in the Supplemental Methods. Fast wide-field Ca^2+^ imaging used an Olympus BX51WI upright microscope equipped with an Evolve 512 EMCCD camera. For mechanical stimulation, an individual neutrophil was gently approached with a 1 μm glass-pipette tip, under visual control, until a deflection in the pipette resistance reported membrane contact.

Glutamate release was first monitored with glutamate-sensitive electrochemical microelectrode biosensors, with a 7 μm tip ^21^ (Sarissa Biomedical Ltd., Coventry, UK); biosensors were calibrated prior to each experiment using 10 μM glutamate application. To visualise glutamate released by individual neutrophils, we used an enzymatic assay as described earlier ^30,31^, with some modifications (Supplemental Methods). Again, glutamate-sensitive fluorescence was collected with a fast-speed Evolve 512 EMCCD camera (Photometrics), at 30-33°C in zero-Mg^2+^ solution.

The statistical analysis accounted for the experimental design involving real-time recordings with direct in situ (same cell) comparison, independent factors, and no longitudinal trials. Correspondingly; independent- and paired-sample *t*-test (Gaussian distribution) or non-parametric tests (Gaussian distribution rejected by Shapiro test) were employed.

## RESULTS AND DISCUSSION

First, to probe directly the function of NMDARs expressed in neutrophils ^21^, we held individual cells in whole-cell mode (current clamp; 0-Mg^2+^ solution). A one-second pulse of NMDA (100 μM) and the NMDAR co-agonist glycine (1 mM) using a rapid-solution-exchange system (Fig. 1A) ^23^ evoked inward current, which was blocked either by extracellular Mg^2+^ (2 mM), by the specific NMDAR antagonist APV (50 μM), or by the selective antagonist of NR2B-containing NMDARs Co101244 (1 μM, Fig. 1B-C). Similarly, it evoked a prominent Ca^2+^ rise detected using the Ca^2+^ indicator Fluo-4 (Fig. 1D), which was, again, blocked by Mg^2+^ or Co101244 (Fig. 1E-F). Towards the end of such tests, we confirmed the cell Ca^2+^ signalling capacity by applying the protein kinase C activator phorbol myristate acetate (PMA, 1 μM), which triggered a further Ca^2+^ elevation (Fig. 1E-F). These observations provide direct electrophysiological evidence for functional NMDARs in neutrophils, predominantly GluN2B subtype, revealing an important mechanism of Ca^2+^ mobilisation in these cells. However, neutrophil depolarisation could also produce a robust Ca^2+^ rise that was insensitive to APV (Fig. 1H), confirming the additional, voltage-dependent and NMDAR-independent routes of Ca^2+^ entry, such as the voltage-gated Ca^2+^ channel Kv1.3 ^32^.

**Figure 1.**
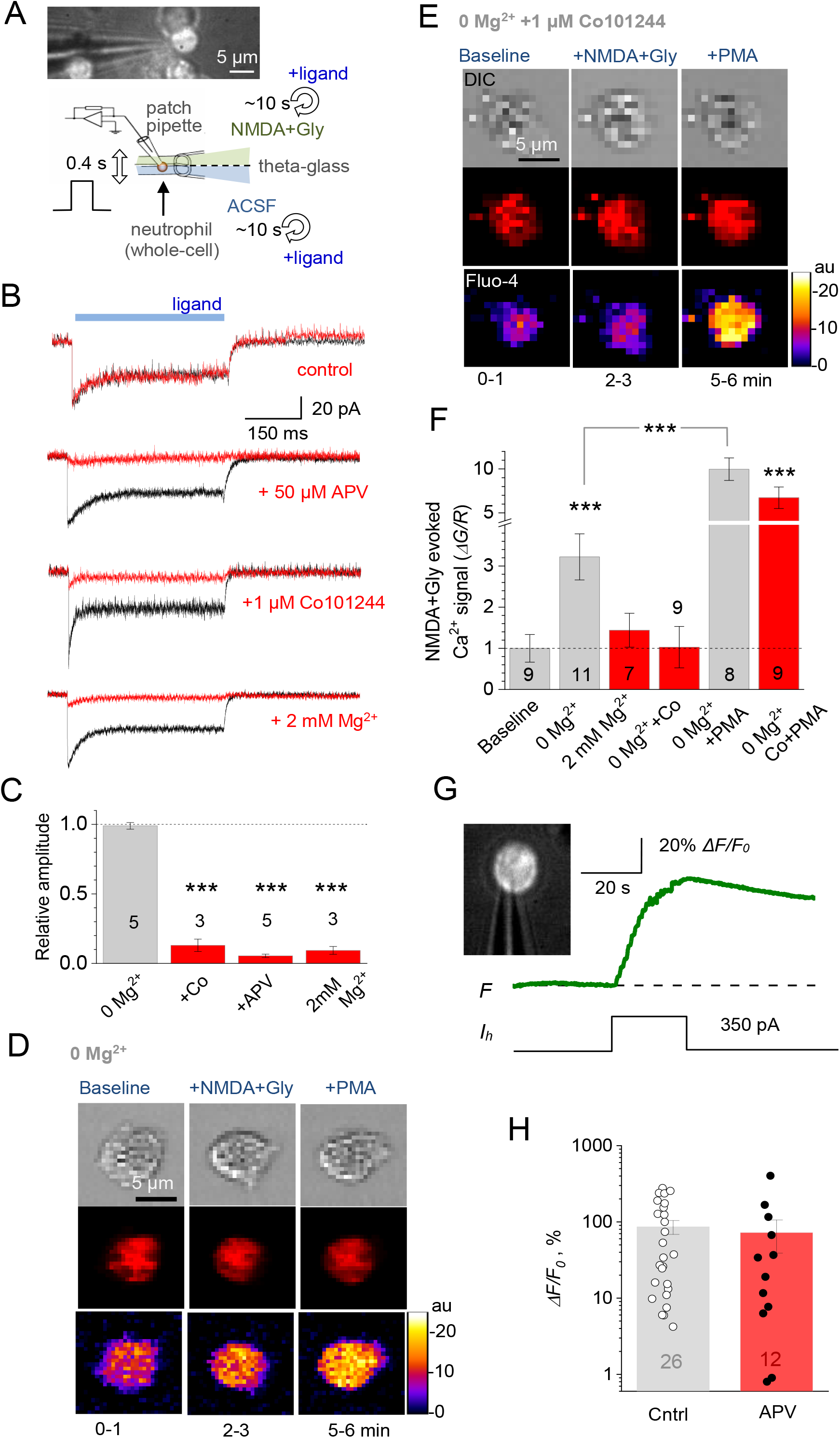
Activation of NMDA receptors in human neutrophils. (A) Diagram depicting probing of NMDARs with a rapid-exchange system [28]: a neutrophil (held in whole-cell) is stimulated pharmacologically by applying different solutions through two channels of ϴ-glass pipette (tip diameter ~200 μm) mounted on a piezo-drive to enable the ultra-fast delivery (<1 ms resolution; solutions in each channel exchanged within 10 s). (B) Representative whole-cell currents (patch pipette 4-7 MΩ, 1-2 μM tip) recorded in human neutrophils in response to locally applied NMDA (1 mM) and glycine (1 mM), in zero Mg^2+^, 2 mM Mg^2+^, and in the presence of NMDAR antagonists APV (50 μM) and Co101244 (1 μM) in zero Mg^2+^ (control), as indicated; same-cell pharmacological manipulations applied at ~20 s intervals. (C) Summary of experiments shown in (B); mean ± SEM (amplitude over the 300-500 ms pulse segment), normalised to control (sample size shown). ***p < 0.01. (D) Characteristic images of a neutrophil (grey DIC image, top raw) loaded with CellTracker™ Red (red channel, middle) and Fluo 4-AM (green, bottom), before and after bath application (2-3-min duration) of NMDA (100 μM) and glycine (50 μM), and added PMA (1 μM), as indicated; experimental timing as shown; false colour scale: relative intensity, arbitrary units (au); pixel size ~120 nm (near diffraction limit). (E) Experiment as in (D), but in the presence of the selective NR2B-containing NMDAR antagonist Co101244 (1 μM); other notations as in (D). (F) Statistical summary of experiments shown in (D-E): *ΔG/R*, Ca^2+^ sensitive fluorescent green Fluo-4 signal (*ΔG*) evoked by NMDA+Gly application and normalised to the CellTracker™ Red signal (*R*), to cancel out fluctuations in focus / brightness; bars, mean ± SEM normalized to control (sample size shown); ***p < 0.001 (one-way ANOVA, Bonferroni post-hoc test). (G) One-cell example (neutrophil held in whole-cell, patch pipette is seen) showing that depolarization current induces prominent Ca^2+^ mobilization; inset, DIC+Fluo-4 AM image; traces: *F*, Fluo-4 fluorescence signal; *I*_h_, holding current. (H) Summary of experiments shown in (G): depolarization-induced [Ca^2+^]_in_ rise represented by the Fluo-4 *ΔF/F*_0_ signal; dots, individual cells; bars: mean ± SEM; sample size shown.

Second, to obtain direct evidence for remote neutrophil-neutrophil interaction, we held two neutrophils positioned 5-15 μm apart, in whole-cell mode (Fig. 2A), and applied a depolarising stimulus to one cell to trigger its intracellular Ca^2+^ rise (as Fig. 1G). Remarkably, this evoked an APV-sensitive inward current in the other patched cell (Fig. 2B). The response occurred 70-140 ms after the stimulus (Fig. 2C, left). With glutamate diffusivity of ~0.7 μm^2^/ms ^33^, diffusion theory indeed suggests a 50-150 ms post-release lag, depending on the separating distance, before glutamate concentration reaches its peak (Fig. 2C, right). Intriguingly, for the glutamate concentration to peak at the minimum level sufficient for NMDAR activation (0.5-1 μM) 5-10 μm away, the neutrophil should generate a 5-10 μM glutamate source emanating from its surface (see Supplemental Methods).

**Figure 2.**
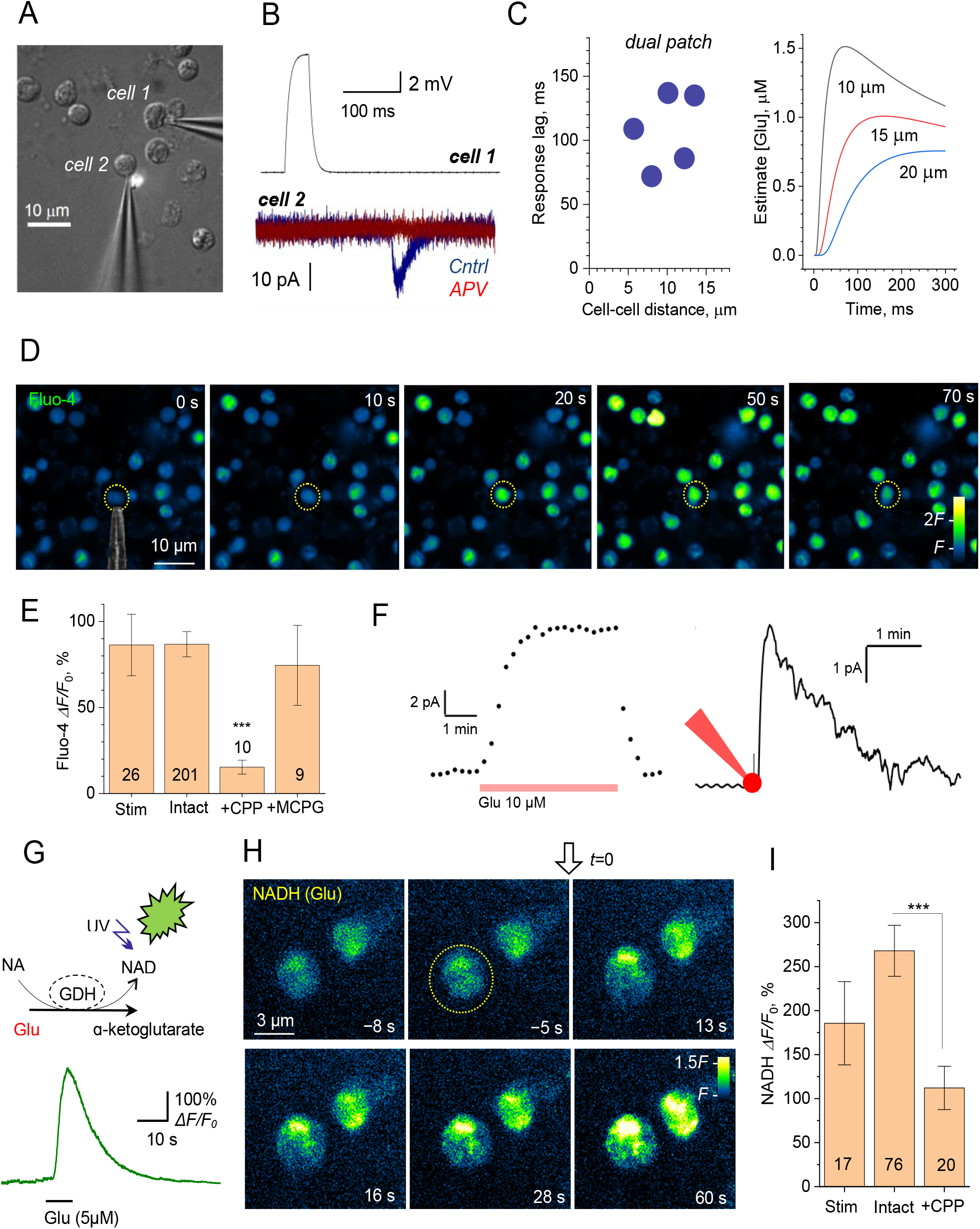
Stimulation of one neutrophil elicits NMDAR-mediated currents, Ca^2+^ rise, and glutamate release in neighbouring neutrophils. (A) Illustration, dual whole-cell recordings from two human neutrophils (cell 1 and cell 2) semi-suspended in culture; patch-pipette tips can be seen; some neutrophils lie destroyed after unsuccessful dual-patch attempts. (B) Example traces from experiments in (A). A 50 ms current pulse applied in cell 1 (upper trace; current clamp) triggers an inward current deflection in cell 2 (voltage clamp), which is blocked by 50 μM APV; bath medium contains 1 mM glycine. (C) Summary of experiments shown in (A-B) for the five recorded neutrophil pairs (*left*), and theoretical estimates of the glutamate concentration time course at three different distances, as indicated, from a small source neutrophil (*right*); see Supplemental Methods for detail. (D) Single-cell induced spread of Ca^2+^ signals among neutrophils. Time-lapse sequence (green Fluo-4 channel) in a semi-suspension of human neutrophils; dotted circle, neutrophil held in whole-cell mode (patch pipette fragment from DIC image shown) enabling one-cell electrical stimulation; electric stimulus (50 pA current) is applied at time zero to the patched cell; false colour scale of Fluo-4 fluorescence *F*, as indicated. (E) Statistical summary of experiments in (D): amplitude of Fluo-4 fluorescence response (mean ± SEM) at the stimulated cell (Stim), neighbouring intact neutrophils (Intact), and in the presence of the NMDAR blocker CPP (10 μM; +CPP) or the wide-range metabotropic glutamate receptor blocker MCPG (400 μM, + MCPG); numbers inside bars, sample size, ***, p < 0.001 (two-sample *t*-test, with respect to the three other sample means). (F) *Left*: Glutamate biosensor sensitivity test: a characteristic response to a 10 μM glutamate application, as indicated; dots, sensor record taken every 15 s. *Right*: A glutamate biosensor record during 10 s whole-cell current injection (red cell-patch diagram) in an individual neutrophil held in whole-cell, as in (D). (G) *Top*: Schematic of the glutamate imaging; A fluorescent enzymatic-based assay involves conversion of β-nicotinamide adenine dinucleotide (NAD) by L-glutamic dehydrogenase (GDH) (in the presence of glutamate) to NADH that fluoresces upon UV light excitation. *Bottom*: Assay sensitivity text; a fluorescence response to a pressure pulse of 5 μM glutamate (~1 μm pipette tip; fluorescence intensity average over a 10μm x10 μm area around the tip). (H) Time-lapse series of fluorescence images (NADH channel) depicting a prominent rise in extracellular glutamate near the two neutrophils upon mechanical stimulation (light touch by a 1 μm micropipette tip) of one cell (dotted circle) at *t* = 0 s, as indicated; false colour scale; note that the glutamate concentration signal drops sharply away from the cell surfaces reflecting rapid dilution in the bath medium. (I) Statistical summary of experiments in (H): amplitude of glutamate-sensitive NADH fluorescence response (mean ± SEM) at the stimulated cell (Stim), neighbouring intact neutrophils (Intact), and in the presence of the NMDAR blocker CPP (10 μM; +CPP); numbers inside bars, sample size; ***, p < 0.001 (two-sample *t*-test).

To understand whether this signal exchange triggers intracellular Ca^2+^ mobilisation, we monitored intracellular Ca^2+^ in pairs or groups of neutrophils. The depolarising stimulus applied to one cell triggered a transient Ca^2+^ rise in its neighbours (Fig. 2D), which was blocked by the specific NMDAR antagonist CPP but not by the broad-range metabotropic receptor antagonist MCPG (Fig. 2E).

Finally, we sought to understand if glutamate released from one neutrophil prompts glutamate release from its neighbours. We first used a glutamate-specific biosensor ^34^ (Fig. 2F, left) to show that a depolarising stimulus at one cell produces a micromolar-range glutamate response at 5-15 μm from it (Fig. 2F, right), as suggested previously ^20,21^. Next, to visualise directly the fate of glutamate during neutrophil signalling, we employed an established enzymatic assay for extracellular glutamate imaging ^30,31,35^ (Fig. 2G, top), also confirming its robust glutamate sensitivity (Fig. 2G, bottom). We thus monitored pairs or small groups of neutrophils using the enzymatic assay and found that even a brief stimulation of one neutrophil (light touch of a patch-pipette) induced pronounced glutamate rises, both at the stimulated and the neighbouring cell (Fig. 2H). Relating these data to the glutamate-induced NADH signal (Fig. 2G, bottom) suggests that the endogenously released glutamate could indeed reach a local level of several micromolar, near the surface of both stimulated (n = 17) and remote cell (n = 76; p = 0.362; Fig. 2I). This level appears not far from the concentration range of ~8 μM considered optimal for neutrophil chemotaxis and cytoskeleton polarization ^36^. Intriguingly, blocking NMDARs with CPP suppressed glutamate-induced glutamate release incompletely (Fig. 2I), which could be the effect of the residual glutamate diffusing from the stimulated cell in an NMDAR-independent manner.

Cooperative neutrophil behaviour, such as swarming, appears to rely on Ca^2+^-induced release of chemo-attractants, with the paracrine-autocrine actions providing positive-feedback signal amplification ^13–15^. This involves release of various signalling molecules that target their receptors on the neutrophil surface ^11,12^. Previous studies have indicated that neutrophils express NMDARs and can release glutamate ^20,21^ while also operating powerful glutamate uptake ^37^. We therefore hypothesised that glutamate-induced glutamate release could contribute to the self-propagating molecular signalling among neutrophils. Our experiments demonstrated direct signal exchange among human neutrophils: this involves Ca^2+^ dependent glutamate release from a one cell, which activates NMDARs in the neighbouring neutrophil(s), which in turn induces Ca^2+^ entry in the receiving cells prompting them to release glutamate. This signal propagation cycle could be therefore regenerative. Interesting, NMDAR activation requires membrane depolarisation to relieve the Mg^2+^ block ^38^ whereas neutrophils appear to undergo drastic depolarisation during their activation ^39^. The latter suggests that our findings could be particularly relevant to neutrophil behaviour under activation.

## Author Contributions

SS initiated the study and carried out patch-clamp experiments and analyses in individual cells; LB carried out imaging tests in individual cells; PM did initial biochemical probing; AGA and GLA helped with neutrophil preparations; AVG provided glutamate and D-serine sensor data; OK carried out imaging and patch-clamp experiments and analyses in neutrophil ensembles and wrote experimental parts of the manuscript draft; DAR narrated the study, carried out some image and data analyses and completed the manuscript draft.

## Institutional Review Board Statement

University College London Ethics Committee approval No 5779/001.

## Informed Consent Statement

Informed consent was obtained from all subjects involved in the study.

## Data Availability

The data presented in this study are available on request from the corresponding author. The data are not publicly available due to ethical concerns.

## SUPPLEMENTAL MATERIALS AND METHODS

### Preparation of human neutrophils

Neutrophils were isolated from blood samples obtained by venepuncture from healthy volunteers according to protocols approved by the UCL Research Ethics Committee (UCL Queen Square Institute of Neurology, London, UK), with the informed consent of participants.

For the isolation of neutrophils, we used a method of dextran sedimentation and differential centrifugation through a Ficoll-Hypaque density gradient as described in detail elsewhere {Park, 2002 #24}. Briefly, a sample of the whole blood was suspended in sodium citrate solution, and a suspension of neutrophils was obtained by sedimentation in 2% dextran (Sigma, UK) in 0.9% NaCl for 45-60 min at room temperature. The neutrophil-enriched upper layers of the suspension were collected and centrifuged (1150 rpm, 10 min at 4 °C). Residual erythrocytes were removed by hypotonic lysis, and the obtained suspension was further centrifuged (1300 rpm, 6 min at 4 °C); the pellet was further re-suspended in PBS and purified by gradient centrifugation over Ficoll-Hypaque (Sigma, cat. 1077; 1500 rpm, 30 min at 4 °C). The resulting pellet containing neutrophils was finally re-suspended in HBSS (0-Mg^2+^/0-Ca^2+^); cells were plated on coverslips coated with poly-DL-Lysine (1 mg/ml) and laminine (20 μg/ml) and were maintained until used (37°C, 5% CO_2_). With this protocol, neutrophils remained lightly attached to the coverslip, forming a semi-suspension: this partly restricted their movement while allowing for patch-clamp and fluorescence imaging experiments in individual cells. At the same time, the cells were not flattened by strong adhesion and showed considerable morphological plasticity on the microscopic scale. We identified healthy cells by their normal morphology, including pseudopodia motility, stimulus evoked transient Ca^2+^ responses, normal NMDAR-dependent activity, and evoked glutamate release. The cultures still contained a significant proportion of healthy, stimulation-responsive neutrophils for up to 24 h, but the present data were collected within the fist few hours post-isolations.

### Patch-clamp electrophysiology

Visualised patch-clamp recordings from the neutrophils were performed using a Multipatch 700B amplifier controlled by pClamp 10.2 software package (Molecular Devices, USA). For the recordings, cells on a glass coverslip were placed in a recording chamber mounted on the stage of an Olympus BX51WI upright microscope (Olympus, Japan). The perfusion medium contained (mM): 119 NaCl, 2.5 KCl, 1.3 MgSO_4_, 2.5 CaCl_2_, 26.2 NaHCO_3_, 1 NaH_2_PO_4_, 10 glucose and was continuously bubbled with 95% O_2_ and 5% CO_2_ (pH 7.4; 290–298 mOsm).). For the recordings of the NMDAR-mediated currents, the medium was changed for a modified zero-Mg^2+^ solution that contained (mM) 119 NaCl, 2.5 KCl, 2.5 CaCl2, 26.2 NaHCO3, 1 NaH2PO4, 23 glucose (pH 7.4; 290–298 mOsm). The NMDAR-mediated currents were measured in voltage-clamp mode, V_hold_ = −70 mV at 33-34°C. Signals were digitized at 10 kHz. Patch pipettes were pulled from borosilicate glass and fire-polished to 4-7 MΩ (1.5-3 μm tip). The intracellular pipette solution for voltage-clamp experiments contained (mM): 120.5 CsCl, 10 KOH-HEPES, 2 EGTA, 8 NaCl, 5 QX-314 Br^−^, 2 Mg-ATP, 0.3 Na-GTP, 10 Na-phosphocreatine, pH and osmolarity adjusted to 7.2 and 295 mOsm, respectively. For whole-cell experiments in current mode, an internal solution contained (in mM) 135 K-methylsulphonate, 4 MgCl_2_, 10 Tris-phosphocreatine, 10 HEPES, 4 MgATP, 0.4 GTP-Na (pH 7.2, osmolarity 290 mOsm, pipette resistance 4–7 MΩ). Stimulating protocol consisted of a series of 500-ms depolarizing rectangular current pulses injected into a neutrophil with an increment of 20 to 30 pA (input current was injected up to 300-350 pA).

The previously established fast-application system {Sylantyev, 2013 #9745} included a theta-glass application pipette with ~200 μm tip diameter attached to the PL127.11 piezo actuator driven by the E-650.00 LVPZT amplifier (both from PI Electronics). Routinely, one pipette channel was filled with the bath solution and the other channel was filled with the bath solution containing 100 μM NMDA and 1 mM glycine, or alternatively with NMDA and glycine, plus NMDAR modulators. The pressure in the application pipette was regulated by a two-channel PDES-02DX pneumatic micro ejector (npi electronic GmbH) using compressed nitrogen. To test the effects of various ligands, the application solutions in both theta-glass pipette channels could be exchanged within 10-12 s during the experiment using dedicated pressurised micro-circuits. NMDAR ligands were applied in 400 ms pulses five seconds apart.

### Diffusion time course estimates

The glutamate concentration time course *C*(*r*,*t*) at distance *r* from the glutamate-releasing cell was estimated, as a first approximation, using the classical equation for a small source in an infinite volume:

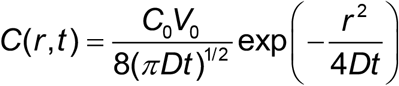

where *D* = 0.7 μm^2^/ms is glutamate diffusivity, and *C*_0_ stands for glutamate concentration at *t* = 0 within small source volume V_0_. For the sake of simplicity, in the calculations we assumed that that volume from which glutamate is released can be represented by a 0.5 μm layer around a 6 μm wide spherical neutrophil. To reach the minimum glutamate concentration required to activate NMDARs (0.5-1 μM), this equation predicts *C*_0_ in the range of ~5 μM (or equivalently, ~10 μM within a 0.25 μM layer around the neutrophil source).

### Two-photon excitation (2PE) fluorescence imaging

For live-cell imaging, neutrophils were loaded with a morphological tracer CellTracker™ Red (5 μM, Invitrogen) and Ca^2+^-indicator Fluo-4/AM (5 μM, Invitrogen) in the presence of Pluronic F-127 (0.02%, Invitrogen) for 10 minutes at 30°C. After incubation with the dyes, cells were washed out for 10 minutes in a medium containing (mM): 119 NaCl, 2.5 KCl, 1.3 MgSO_4_, 2 CaCl_2_, 26 NaHCO_3_, 1.25 NaH_2_PO_4_, 12 glucose (95% O_2_ and 5% CO_2_; pH 7.4, 290–300 mOsm). Imaging was performed in a medium of the same composition containing either 0 or 2 mM MgSO_4_, as specified, at 30-33°C. Imaging was carried out using an Olympus FV-1000MPE system optically linked to a Ti:Sapphire MaiTai femtosecond pulse laser (SpectraPhysics Newport) at λ^2P^_ex_ = 800 nm, with appropriate emission filters (Fluo-4: 515-560 nm band; Cell tracker red: 590-650 nm band), as detailed previously {Jensen, 2021 #10669;Kopach, 2022 #11024}. For the time-lapse recordings, *z*-stacks of fluorescent images (containing 5-10 cells within the field of view) were collected in a 1-min increment for 2 min before (baseline) and upon application of NMDA (100 μM) and glycine (50 μM) (bath application for the next ~3 minutes). At the end of each experiment, the protein kinase C activator phorbol myristate acetate (PMA, 1 μM) was added for 2 min as a functional test. For the analysis, the Fluo-4 signal (*G*, green channel) was normalised to the Cell tracker fluorescence (*R*, red channel), and changes in Ca^2+^ level were expressed as *ΔG/R* after background subtraction. Only cells displaying a stable baseline and robust (>two-fold) *ΔG/R* increase in response to PMA, with no Fluo-4 saturation, were included in the statistics.

### Fast wide-field fluorescence imaging

Neutrophils were loaded with Fluo-4 AM (5 μM, Invitrogen) in the presence of Pluronic F-127 (0.02%, Invitrogen) for 15 minutes at 37°C. After loading, cells were washed out for approximately 30 minutes in a Ringer solution containing (in mM) 126 NaCl, 3 KCl, 2 MgSO_4_, 2 CaCl_2_, 26 NaHCO_3_, 1.25 NaH_2_PO_4_, 10 glucose, equilibrated with 95% O_2_ and 5% CO_2_ (pH 7.4; 290–300 mOsm). Experiments were performed at 30-33°C, using an Olympus BX51WI upright microscope (Olympus, Japan) equipped with a LUMPlanFI/IR 40×0.8 objective and an Evolve 512 EMCCD camera (Photometrics). A source of fluorescent light was an X-Cite Intelli lamp (Lumen Dynamics). For the time-lapse recordings, 700 to 3000 images were acquired in the stream-acquisition mode (exposure times from 20 to 100 ms), using MetaFluo (Cairn Research Ltd, UK) or Micromanager 4.1 (freely available ImageJ plugin) software. To collect images at high resolution, various digital zooms were used (100–200 nm per pixel). Changes in the intracellular Ca^2+^ level were expressed as the changes in Fluo-4 fluorescent signal over the baseline (*ΔF/F_0_*) after subtraction of background fluorescence and correction for bleaching.

For mechanical stimulation, an individual cell was gently tapped with a glass pipette over the cell surface, under visual control, while monitoring changes in the pipette resistance (confirming the contact with the cell membrane).

### Glutamate recordings with an electrochemical sensor

For the assessment of changes in the extracellular glutamate level across isolated neutrophils, we used the glutamate-sensitive electrochemical microelectrode biosensors, with a tip of 7-μm in diameter (Sarissa Biomedical Ltd., Coventry, UK). For the recordings, a glutamate biosensor was placed close to the stimulated neutrophil (within 5-15 μm), with a null biosensor located apart. After positioning, the basal current was recorded for a few minutes before (baseline) and upon cell stimulation. Biosensors were calibrated prior to each experiment using 10 μM of glutamate.

### Glutamate imaging

Fast wide-field imaging was used to visualise glutamate released by individual neutrophils in combination with an enzymatic assay as described earlier {Bezzi, 1998 #2074;Ayoub, 1998 #26}, with some modifications. Before the recordings, isolated neutrophils were perfused with a HEPES-based medium containing (in mM) 135 NaCl, 5 KCl, 2 CaCl_2_, 2 MgCl_2_, 10 HEPES, 10 glucose (pH 7.4; 290–300 mOsm). The bath medium was supplemented with L-glutamic dehydrogenase (GDH, 60 U/ml, Sigma, UK) and β-nicotinamide adenine dinucleotide (NAD, 1 mM). In the presence of glutamate, L-glutamic dehydrogenase (GDH) reduces NAD(+) to NADH, a product that fluoresces when excited with either UV light or in two-photon excitation mode {Innocenti, 2000 #10753}. The detection of released glutamate thus relies on the NADH-mediated fluorescence following the GDH-catalyzed conversion of NAD to NADH in the presence of glutamate. Imaging was performed in a semi-suspension of isolated neutrophils using an Olympus BX51WI upright microscope (Olympus, Japan) equipped with a LUMPlanFI/IR 40×0.8 objective and appropriate emission filters (U-MWU2 filter set, Olympus). Fluorescence was collected with a fast-speed ultrasensitive Evolve 512 EMCCD camera (Photometrics). Glutamate imaging recordings were performed at 30-33°C in zero-Mg^2+^ solution, with no perfusion running. Experiments were performed at 30-33°C. Changes in the NADH fluorescence were calculated as ΔF/F_0_ after the subtraction of background fluorescence. Background subtraction was performed for the time-lapse sequence, by subtracting the optical image before stimulation throughout *T*-stacks to eliminate basal NADH fluorescence and the contaminating fluorescence, such as autofluorescence of enzymes.

### Statistical analysis

The experimental design involved real-time recordings with direct in- situ (same cell) comparison, independent factors, and no longitudinal trials; independent- and paired-sample *t*-test (Gaussian distribution) or non-parametric tests (Gaussian distribution rejected by Shapiro test) were used, as required.

